# Combined inference of known and novel mutational signatures with ReDeNovo

**DOI:** 10.64898/2026.02.05.703798

**Authors:** Ziynet Nesibe Kesimoglu, Ermin Hodzic, Jan Hoinka, Bayarbaatar Amgalan, M.G. Hirsch, Teresa M. Przytycka

## Abstract

Mutational signatures represent characteristic mutational patterns imprinted on the genome by mutagenic processes. They can provide information about the impact of the environmental and endogenous cellular processes on tumor mutations and can suggest treatment. Analysis of presence and strength of mutational signatures in cancer genomes has become a cornerstone in analysis of new and legacy cancer data. However, a precise inference of novel (*de novo*) signatures requires a large set of genomes, and methods focusing on estimating the presence of previously defined signatures are unable to uncover potential novel signatures that might emerge in new data. Thus, reliable methods to address these challenges are needed. We formally define the Combined Mutational Signature Inference Problem (CMSI) for the identification of known signatures and the inference of novel signatures in cancer data. CMSI represents non-convex optimization, and we provide a heuristic algorithm, ReDeNovo, to solve it efficiently. We extensively validated ReDeNovo on simulated data, evaluating its ability to precisely estimate presence and exposure to known signatures and to discover of novel signatures. On both tasks ReDeNovo outperformed existing approaches. In real biological data, ReDeNovo identified signatures missed by previous analyses and defined a new signature related to UV light exposure. ReDeNovo method provides a new and powerful tool to uncover mutational signatures. ReDeNovo is available from https://github.com/ncbi/redenovo.

## 1 Introduction

DNA molecules in the cells of living organisms are subjected to various mutagenic processes leading to the accumulation of mutations. These processes can be endogenous or exogenous, such as background stochastic DNA damage and repair, cancer-related aberrations of the DNA maintenance machinery, and mutations triggered by carcinogenic exposures, such as smoking. Most acquired mutations are benign, yet they offer valuable insights into the mutational processes that generated them. *Mutational signatures* are a representation of characteristic mutational patterns that different mutagenic processes imprint on the genome [1, 2, 3] and provide a link between mutagenic processes and somatic changes (though not all signatures have a known etiology). In particular, they provide a way to detect environmental exposures that contribute to carcinogenesis [4]. The presence of specific signatures can be predictive of prognosis and response to therapy [5, 6]. For example, patients with Homologous Recombination Deficiency (HRD) benefit from PARP inhibitor therapy and thus the presence of an HRD signature can be used as its marker [7, 8].

Mutational signatures are most commonly defined as a multinomial distribution over a set of *mutation categories*. For single base substitutions (SBS), the classical definition of mutation categories is based on the mutational change and the trinucleotide context in which the mutation occurs, yielding 96 mutation categories [1, 3]. Following the seminal work of Alexandrov et al. [2], the fundamental assumption common to most papers is that mutagenic processes are additive, implying the mutations in each individual’s genome can be represented as a linear combination of mutations attributed to individual signatures, where the number of mutations attributed to a given signature is called its *exposure* or *activity*. While non-additive models are beginning to emerge [9, 10, 11], the additivity assumption remains the most frequently used. Under this assumption, many mutational signatures have been identified and analyzed, providing a convenient and informative reference for further studies, including the study of the dependence of signature presence on local genomic context (e.g. [12, 13, 14]), inference of causality [15, 16], or for developing supervised machine learning models [17, 18]. Databases of mutational signatures such as COSMIC [19] or the Compendium of Mutational Signatures of Environmental Agents [20] collect information related to signatures and their proposed links to specific mutagenic processes. The evidence that links signatures to mutagenic causes varies: some are supported by mechanistic explanations, some by associations with potential causes, while the etiology of many others remains unknown.

New cancer studies routinely include analysis of the presence and activity of mutational signatures in the cohort analyzed. Approaches for identifying these signatures in tumor data can be broadly classified into two groups: *de novo* methods and *refitting* methods. *De novo* methods aim to uncover mutational signatures and their corresponding activities in new datasets without leveraging the knowledge about previously defined signatures. However, a confident *de novo* estimation of signatures requires a very large set of genomes. This challenge is bypassed by refitting methods. These methods are tailored to uncover only known, previously-defined, signatures and their activities. While computationally more tractable, refitting methods cannot uncover new signatures, which is often an important question in the analysis of new cohorts. This motivates the need for approaches that combine the benefits of both strategies. Such approaches should attempt to explain mutation data using a given set of known signatures, as well as allow for the inclusion of new signatures if the known ones cannot sufficiently explain the data. Recently, the first heuristic approaches that integrate refitting and *de novo* inference have begun to emerge, such as CaMuS [21] or MuSiCal [22]. These methods first acquire candidate signatures via NMF-based optimization, attempt to map them to known signatures, and ultimately select the final list of detected known and novel signatures. Since these NMF-based methods are typically implemented as greedy heuristics, we hypothesize that such front-loaded NMF optimization might often yield inaccurate results. We suggest an alternative framework in which stability and quality control is a core segment of the iterative signature inference process.

In this paper, we define the Combined Mutational Signature Inference Problem and present an algorithm, ReDe-Novo, that provides a heuristic solution to this problem. In contrast to above-mentioned approaches, ReDeNovo infers known and novel signatures dynamically, ensuring that a signature is selected only when the data has abundant support for it, minimizing potential negative impact on the rest of the inference. Using extensive, carefully-designed simulations to reflect properties of real data, we validate our hypothesis and show that this approach provides a better estimation of signatures presence and activity for both predefined and novel signatures. Application of ReDeNovo to cancer patient data further revealed a novel mutational signature with a putative association to UV light.

## 2 Methods

Let *m* be the number of different mutation categories used to define mutational signatures. A *mutational signature* is an *m*-dimensional non-negative vector that represents a probability distribution of the *m* mutations.

For a collection of *n* samples, let *M*_*n*×*m*_ be the matrix of observed counts of the *m* mutation categories across the *n* samples. Assume that we know all ground truth mutational signatures that are active in the given samples: we label them *s*_1_, *s*_2_, …, *s*_*l*_. Then, under the *linear composition model, M* can be decomposed as *M* = *A*_*n*×*l*_ ×*P*_*l*×*m*_ +*ξ*_*n*×*m*_, where i) the rows of *P* are the *l* mutational signatures *s*_1_, *s*_2_, …, *s*_*l*_, and ii) *A* is a non-negative matrix of signature activities in the *n* samples, and *A* is selected to minimize the squared Frobenius norm of *ξ*.

In practice, the ground truth list of exclusively involved mutagenic processes, and thus the corresponding matrix *P*, is often unknown. In *de novo* approaches to mutational signature inference, *M* is decomposed into *A* × *P* through variations of the *non-negative matrix factorization* (NMF) method, without any reference to already known catalogues of mutational signatures. In contrast, *refitting* approaches assume that *P* is the matrix of known mutational signatures, and then compute the corresponding activity matrix *A*. However, recognizing that the ground truth decomposition of *M* is likely to contain some already known mutational signatures, as well as possibly some novel ones, methods increasingly leverage both approaches.

### 2.1 Combined mutational signature inference

Let 𝒞 be the set of all known mutational signatures built from *m* given mutation categories. Let *f* be some objective function used to decompose the observed mutation count matrix *M* into a product of an activity matrix *A* and a mutational signature matrix *P*. For given parameters *δ, ϕ, γ* ∈ (0, 1], we define a restriction 𝒮_*δ,ϕ,γ*_ of the space of all pairs of non-negative matrices that are possible solutions to the decomposition of *M*, such that every pair of matrices (*A, P*) ∈ 𝒮_*δ,ϕ,γ*_ is subject to the following constraints:

1. For every mutational signature row *P*_*k*_, it is either an exact copy of a known mutational signature from 𝒞, or its largest cosine similarity with any element of 𝒞 is smaller than *δ*:

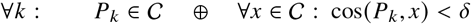
2. Every activity matrix column *A*_*k*_ has at least *ϕ* fraction of samples whose relative activity fraction (relative to the total sum of activities across all samples) is at least *γ*:

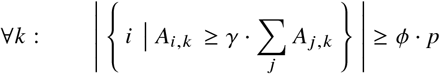

Constraint (1) achieves a combination of *de novo* and refitting approaches by requiring each row of the inferred signature matrix to either be a previously known mutational signature, or to be sufficiently dissimilar from each of them. Constraint (2) aims to avoid inference of signatures if their activity is insufficient and thus difficult to distinguish from noise.

𝒮_*δ,ϕ,γ*_ describes the space of all feasible solutions to the combined mutational signature inference problem for that particular choice of parameters *δ, ϕ, γ*. From that space, we are looking for the solution that is optimal with respect to two competing objectives: minimization of the error of the model, and minimization of the number of signatures used.

#### The Combined Mutational Signature Inference Problem (CMSI)

Given a set of known mutational signatures 𝒞, an observed mutation count matrix *M*, real numbers *δ, ϕ, γ* ∈ (0, 1], and a multiobjective linear scalarization parameter α ∈ ℝ, compute the pair of matrices (*A, P*) ∈ 𝒮_*δ,ϕ,γ*_ which minimizes the multi-objective function 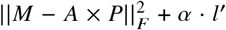, where *l*^′^ is the num-ber of mutational signatures in *P* that are not from 𝒞.

Similar to the NMF problem, the CMSI problem represents non-convex optimization, and there is no known algorithm to find its global optimum in polynomial time.

### 2.2 ReDeNovo

ReDeNovo provides a heuristic solution to the CMSI problem. Assuming certain default values for parameters *δ, ϕ, σ* (which can be changed by the user), ReDeNovo takes as its input the observed mutation count matrix *M* and a reference set of known mutational signatures 𝒞, and decomposes *M* into the activity and signature matrices *A* and *P* in accordance with the objective of the CMSI problem formulation and the *δ, ϕ, σ*-based constraints. ReDeNovo is a two-stage algorithm, consisting of a *recognition* phase followed by a *de novo discovery* phase. The workflow of ReDeNovo is shown in Figure 1 (also see Supplementary Section 2 for gradient descent optimization details and Supplementary Section 3 for algorithm details).

**Figure 1.**
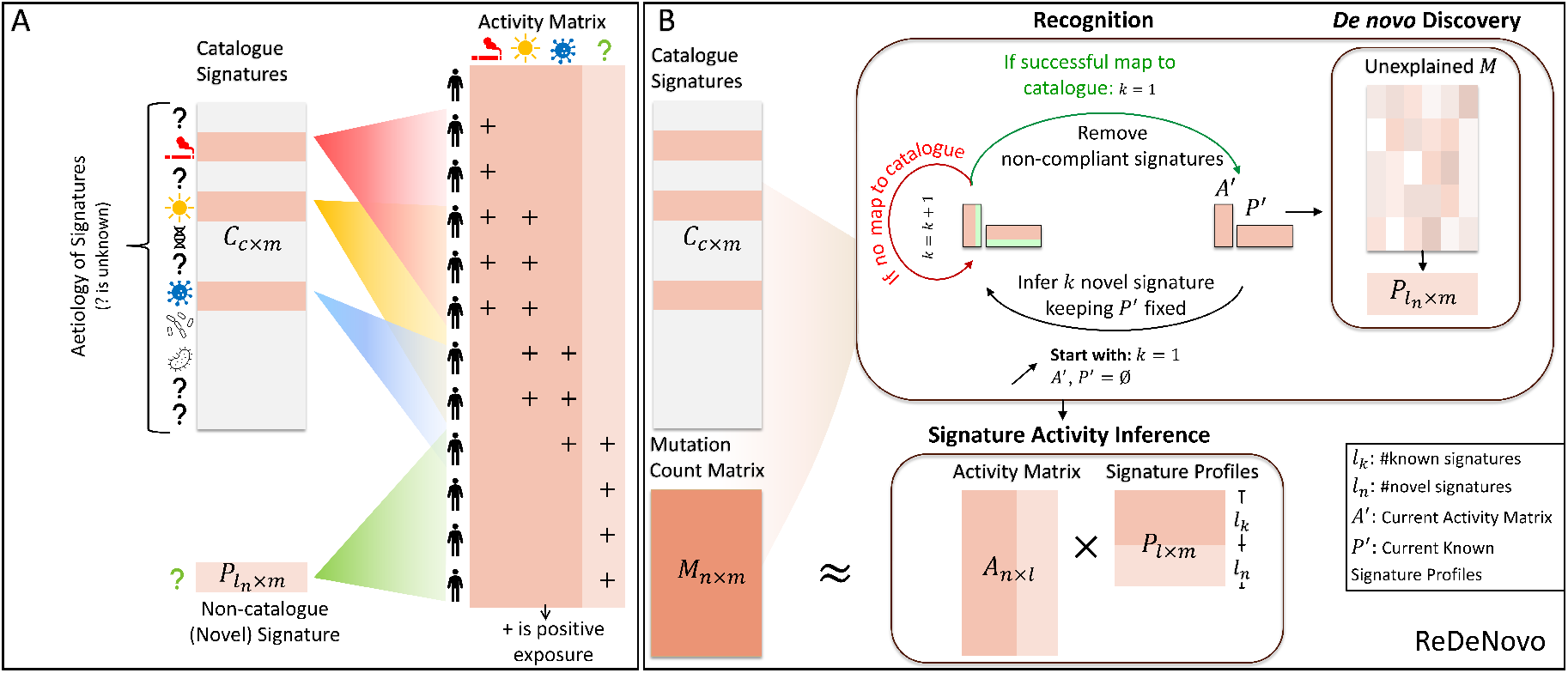
Overview of ReDeNovo. A. Patients are assumed to accumulate mutations attributed to known and unknown mutational signatures. **B**. The recognition phase iteratively adds signatures from a known catalogue, while removing signatures whose activity scores no longer comply with the CMSI constraints. This iterative process ends when no further known signatures can be inferred. The *de novo* discovery phase employs a similar iterative process with the aim to identify novel signatures, dissimilar from any catalogue signature.

The recognition phase is done in an iterative manner. In each iteration, we aim to identify one signature from the catalogue 𝒞 to be added to the resulting signature matrix *P*. To identify each signature, we run unconstrained hybrid NMF optimization in which the signature matrix consists of fixed rows of currently selected catalogue signatures and *de novo* rows from which the next best-fitting catalogue signature is to be identified through the optimization. Starting with a single *de novo* row, we increase the number of *de novo* rows until either at least one of the resulting *de novo* signatures is sufficiently similar to a known catalogue signature (cosine similarity is ≥ *δ*) and satisfies CMSI constraint (2) or a stopping criterion is met. All such catalogue signatures are considered as candidates to be added to the signature matrix *P*, and among them we select the one signature whose cosine similarity to the corresponding *de novo* row is the highest. We add the new signature to *P*, we recalculate the activity matrix *A* and remove any signatures from *P* with activity scores that no longer pass constraint (2), if any. The recognition phase ends when the above process cannot add any more signatures.

*De novo* discovery phase aims to identify novel mutational signatures. Its process is nearly identical to the recognition phase, with the key distinction that a *de novo* signature is added only if its cosine similarity to every catalogue signature is smaller than *δ* (constraint (2) still applies).

## 3 Results

First, we extensively evaluate ReDeNovo on simulated data and compare its performance to previous approaches. Our carefully designed simulations ensure that simulated data is biologically realistic. Next, we apply ReDeNovo to biological data, demonstrating its power to gain new biological insights missed by other approaches.

### 3.1 ReDeNovo outperforms state of the art tools at known mutational signature recognition

We start by systematically analyzing the performance of ReDeNovo and other tools at recognizing known mutational signatures. For this task, we generated 16 synthetic datasets with only known mutational signature presence and their activities (see Supplementary Section 1 for simulation data details). Here we analyze results obtained from synthetic dataset 1 in detail, while the Supplementary Table 3 summarizes consistent results on the remaining 15. As the reference set of mutational signatures, we used COSMIC v3.4. We then assessed the performance of tools both with no noise in the input mutation count data, as well as with increasing levels of random Gaussian noise.

We compared ReDeNovo to SigProfilerAssignment (v0.2.3) [23], SigProfilerExtractor (v1.2.0) [24], MuSiCal [22], sigLASSO (v1.1) [25], and CaMuS [21]. CaMuS requires the user to set two parameters in order to run the method: the expected number of known mutational signatures to recognize, as well as the expected number of novel signatures. For the benefit of the method, we set those parameters to the values consistent with the test data, and we label the results “CaMuS (oracle).” In practice, these parameters are unknown and are inferred by ad hoc strategies.

Since sigLASSO and CaMuS output normalized activity matrices with weights representing the proportional contribution of each signature rather than discrete counts, we rescaled the activities by multiplying each samples’s activity values by the sample’s total mutation burden, to ensure comparability across tools.

We evaluate the different methods’ ability to accurately predict the presence or absence of signatures using F1 scores. To quantify the accuracy of the predicted activities, we calculated the root mean square error (RMSE) between the simulated activity matrix and the output activities. Figure 2A summarizes these results for varying levels of background noise in the input mutation count data (exact F1 scores and RMSE values are reported in Supplementary Table 1 and 2, respectively).

**Figure 2.**
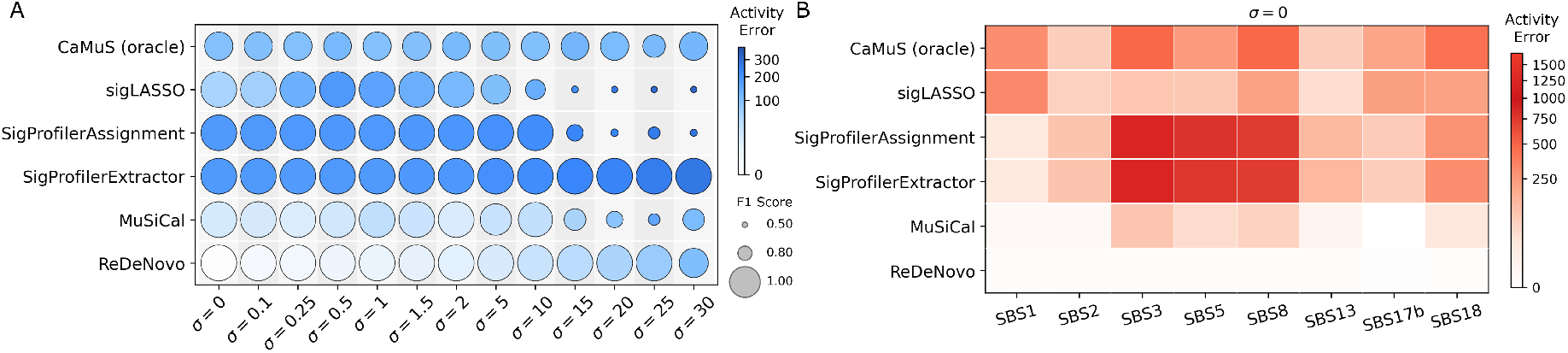
Performance comparison on synthetic data generated using COSMIC mutational signatures. A. Each column represents a different level of standard deviation of Gaussian noise in the input mutation count data. Bubble size shows the F1 score for recovering the ground truth signature set. Bubble color shade shows the RMSE of all the inferred activities. Both reported measures are averaged over 10 runs to account for inherent stochasticity of the tested tools. **B**. Per-signature activity error comparison on synthetic data (*σ* = 0) generated using COSMIC mutation signatures. Root mean square error (RMSE) is calculated for activity estimates of each signature (shown individually in each column), averaged over 10 runs to account for inherent stochasticity of the tested tools. The value shown next to the row (+1) denote the number of additional catalogue signature that the corresponding method mistakenly inferred in at least one run. These represent false positive signatures that were not present in the ground truth signature set.

All tested methods are competitive at inferring the correct signature set for levels of noise *σ <* 10 (Fig 2A). However, across all noise levels, ReDeNovo estimated signature activities more accurately than all other methods. Lower RMSE values of ReDeNovo imply that even though the tools are comparable at the task of recovering the ground truth signature set globally, they underperformed at correctly estimating the signature activity per patient.

At higher noise levels (*σ* ≥ 5), some of the tested methods proved vulnerable to inference of a large number of signatures that are not present in the ground truth set (Figure 2 and Supplementary Figure 2). ReDeNovo and SigProfilerExtractor proved resilient to this, with only a single false positive inferred by ReDeNovo at the highest tested noise setting.

To analyze which signatures are most challenging to be recognized, we evaluate per-signature activity error (RMSE). Figure 2A shows the results for the case of no noise in the input (Figure 2B shows the results for all noise levels). Flat signatures SBS3, SBS5, and SBS8 proved to be the most challenging. ReDeNovo consistently infers the activity of these signatures more accurately than other methods, even as the noise levels are increased (Figure 2 and Supplementary Figure 2).

The results show that ReDeNovo consistently outperformed or matched other tools across all evaluated datasets, including under varying noise levels, underscoring its robustness in accurately identifying active signatures.

### 3.2 ReDeNovo outperforms state of the art tools at *de novo* mutational signature recognition

We evaluated the accuracy of methods at finding *de novo* signature in synthetically generated datasets in which all other signatures are known COSMIC signatures. We focused on the case where there is a single *de novo* signature because the target application of ReDeNovo is for analysis of previously studied cancer types in a new context where we expect a limited occurrence of unknown signatures. Here we focused on synthetic dataset 1, which includes SBS1, SBS2, SBS3, SBS5, SBS8, SBS13, SBS17b, and SBS18 as the fixed set of COSMIC catalogue signatures. We performed a series of tests in which the *de novo* signature was selected from among four different synthesized signatures (see Supplementary Section 1 for simulation details and Supplementary Figure 1 for the profiles of synthesized signatures). Synthetic signatures were generated to have a mutation distribution with multiple peaks, while ensuring that they are different from all COSMIC mutational signatures (cosine similarity ∈ [0.00, 0.71]). In addition, to simulate a broader spectrum of realistic *de novo* signatures, we performed tests in which each individual COSMIC signature in the synthetic dataset was hidden from the COSMIC catalogue given to the methods, and thus turned into a *de novo* signature. We note that SigProfilerExtractor was unable to treat a COSMIC signature as *de novo*, so we could not perform the latter test with this method. To account for inherent stochasticity of the tested tools, we run this test 10 times for each setting, and we report the average values.

ReDeNovo clearly outperformed other methods at the task of *de novo* discovery. ReDeNovo is the only method that consistently recovers the *de novo* signature in every synthetic dataset (Figure 3A). ReDeNovo also has the best average performance on other metrics, including the cosine similarity between the recovered *de novo* signature and ground truth signature, the F1 score for recovering the entire ground true signature set, and the average activity errors (Figure 3, Supplementary Table 4). We note that MuSiCal infers many false positive signatures, both known catalogue signatures, as well as wrong *de novo* signatures (Supplementary Figure 2). This is reflected in MuSiCal’s low ground truth *de novo* signature detection rate (Figure 3A).

**Figure 3.**
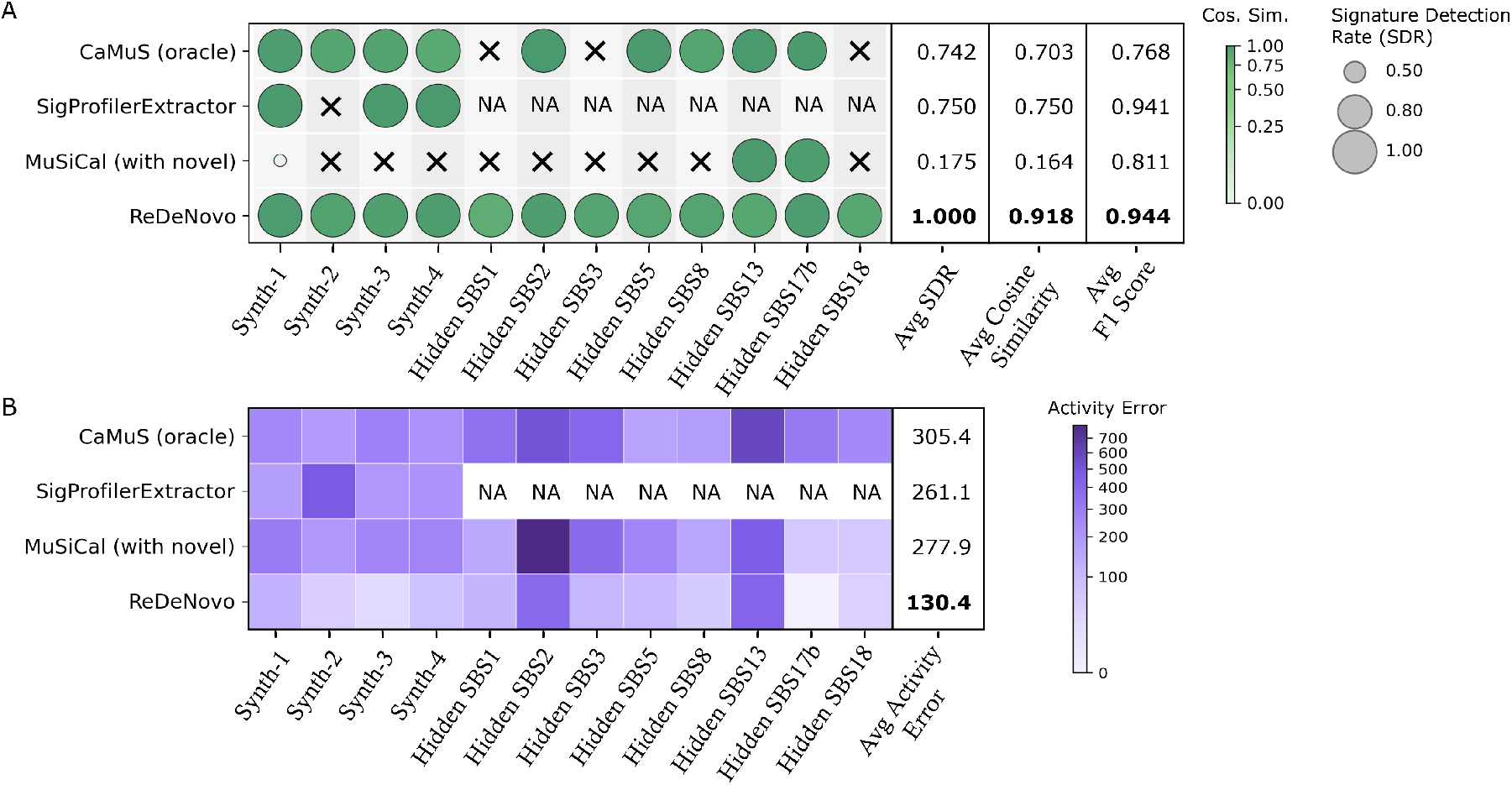
Performance comparison on synthetic data with COSMIC signatures and one *de novo* signature. Each column shows results for a different *de novo* signature in the data, combined with the same fixed set of COSMIC signatures (SBS1, SBS2, SBS3, SBS5, SBS8, SBS13, SBS17b, SBS18). The first four *de novo* signatures are synthetically generated. To make the test as realistic as possible, we also individually hid each COSMIC signature from the input catalogue and treated it as a *de novo* signature. We treat the predicted *do novo* signature as accurately detected if its signature profile has a cosine similarity of at least 0.70 with the ground truth *de novo* signature profile. **A**. Bubble size represents the detection rate of *de novo* signatures. Bubble color shade shows the cosine similarity between real and inferred *de novo* signatures (not inferring any *de novo* signature at all is awarded cosine similarity of 0). **B**. Color shade shows the RMSE of all the inferred activities. All reported measures are averaged over 10 runs to account for inherent stochasticity of the tested tools.

### 3.3 ReDeNovo recognizes previously missed signatures in cancer data

We next applied ReDeNovo to Whole Genome Sequencing (WGS) data for nine cancer types from ICGC-PCAWG [26]. The results are summarized in Figure 4. Because there is no gold standard of truth for biological data, we check if the inferred signatures are consistent with results reported in previous studies for secondary evidence (Supplementary Table 5). We considered previous studies of the same cancer types whether or not the set of patients was the same as in our analysis. Excluding signatures associated with potential sequencing artifacts, 75 out of 83 assignments of COSMIC signatures to cancer types have been previously reported and are consistent with current knowledge (black circles in Figure 4, details in Supplementary Table 4).

**Figure 4.**
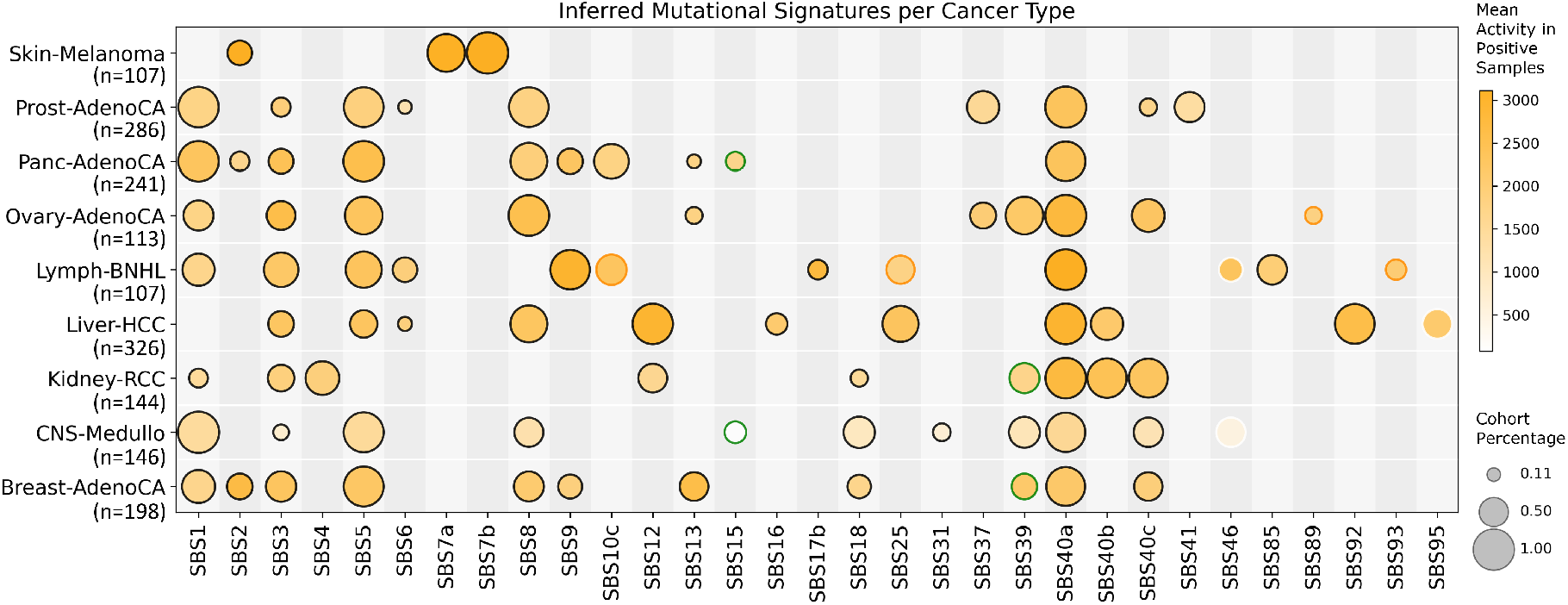
COSMIC mutational signatures recognized by ReDeNovo in ICGC PCAWG WGS cancer datasets. Bubble size represents the percentage of cohort with above the threshold amount of normalized activity, while bubble color represents the mean activity in positive samples. Black outline colors are literature-supported signatures, green outline colors are cases where the signatures were not identified in the literature but there is evidence that the corresponding mutagenic process is likely to be active, white outline colors are signatures corresponding to sequencing artifacts, while orange outline colors are signatures with no supporting evidence found. Sample sizes per cancer type are given under the cancer type labels.

Interestingly, ReDeNovo detected SBS39 in breast and kidney cancers. Although COSMIC currently considers SBS39 to have unknown etiology, recent studies link this signature to two different but related pathways: homologous recombination deficiency (HRD) [27] and Non-Homologous End Joining (NHEJ) [28]. We observe that, in the analyzed dataset, all confirmed assignments of SBS39 were to cancers that also have the homologous recombination deficiency signature, SBS3, assigned to them, supporting the association with mutagenic processes that generate double stands breaks. The novel assignments of SBS39 proposed by ReDeNovo are also assignments to cancers with confirmed presence of SBS3, supporting the correctness of these new assignments. Of the remaining six assignments without direct literature support, two included signatures of unknown etiology (SBS89, SBS93) thus the correctness of their assignment cannot be evaluated based on literature evidence. Two other new assignments are for MMRd signature SBS15 to cancers where there is a literature support for potential presence of MMRd, thus the presence MMRd signatures in a small number of samples as observed in Figure 4 cannot be excluded. However, assignment of SBS15 to CNS MedulloBlastoma is inferred with very small activity, which is atypical for a MMRd signature and thus signaling a potential false positive. There are additional four assignments without an independent support (orange circles in Figure 4). Despite such potential imperfections, nearly all signatures identified by ReDeNovo are supported by previous analyses. We note that we do not evaluate here false negatives as these are hard to resolve based on secondary evidence without a standard of truth. Therefore, for this task, we fully rely on the simulated data presented in the previous section.

### 3.4 ReDeNovo discovers a new UV light-related signature

ReDeNovo predicted only one novel signature in the analyzed datasets. The novel signature is in melanoma and has a single peak for TCC*>*TTC mutations (Figure 5). Historically, the first version of the COSMIC catalogue contained only one signature related to UV light labeled as Signature 7. In the current version (v3.4) of COSMIC, this signature has been replaced by four SBS signatures: SBS7a/7b/7c/7d. Among these, SBS7a and SBS7b contribute disproportionally many mutations to melanoma cancers and were also discovered by ReDeNovo in our melanoma cohort. SBS7a and SBS7b also include a high number of TCC*>*TTC mutations, however, they also feature a high number of other C*>*T mutations: SBS7a also has a high peak for TCA*>*TTA mutations, and SBS7b features a high number of signatures for CCC*>*CTC mutations. The mutations associated with SBS7a have been proposed to be due to UV DNA damage from 6,4-photoproducts while those for SBS7b have been proposed to arise from cyclobutane pyrimidine dimers [29]. The etiology of SBS7c and SBS7d is less understood. These two signatures were not recognized in our analysis of the melanoma cohort under the default parameters of ReDeNovo but could be recognized after removing the influence of 7a and 7b from the mutation count matrix. This is consistent with the fact that the mutational burden attributed to these two signatures is much smaller than the number of mutations attributed to SBS7a/7b and their relative activities do not satisfy the default constraints of ReDeNovo. Subsequent analysis showed that the novel signature co-occurs with previously inferred UV-related signatures. The Pearson correlation coefficient between the cross-patient activity of the novel signature and the activity of SBS7a is 0.83 (Spearman: 0.92) and the correlation coefficient with SBS7b is 0.86 (Spearman: 0.93). This strongly suggests that, similar to SBS7a and SBS7b, this signature is associated with UV DNA damage. The emergence of yet another signature related to UV sun exposure is likely to be a result of both the heterogeneity of the UV wave length exposure and the heterogeneity of the protection mechanisms across human populations. For example, DNA repair efficiency and skin pigmentation play important roles in reducing the mutagenic effect [30] leading to a potential population specific variant of this mutagen.

**Figure 5.**
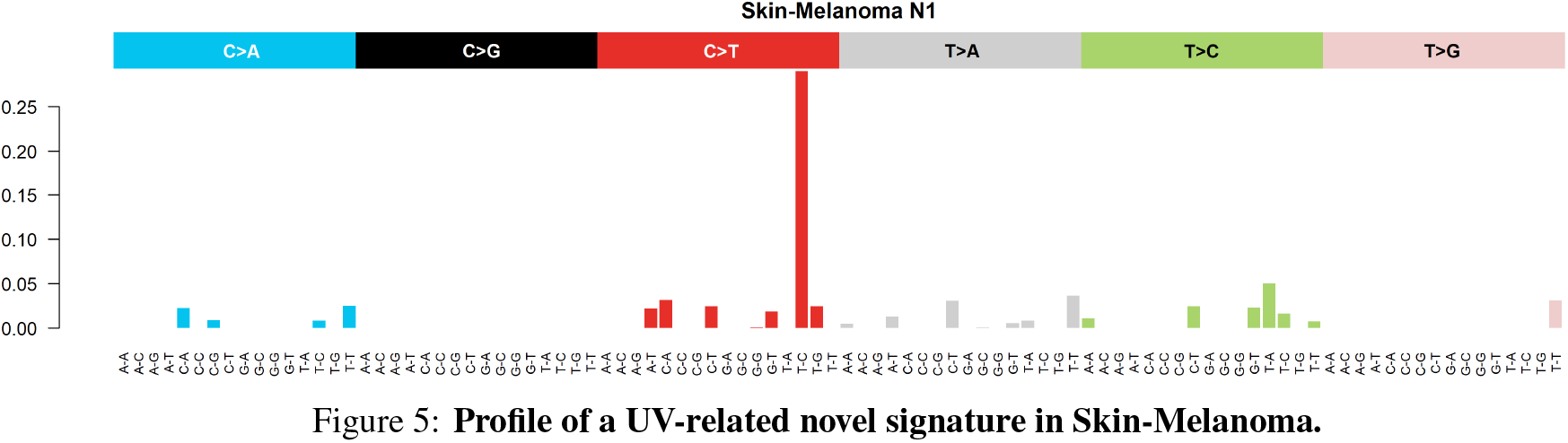
Profile of a UV-related novel signature in Skin-Melanoma.

## 5 Discussion

Analysis of mutational signatures in cancer data is important for understanding the emergence of cancer. It provides insights into mutagenic environmental exposures, deficiency of DNA repair pathways, and other mutagenic processes active in a given cancer. Mutational signature analysis can also help in predicting patient drug response. Over a decade of research has resulted in a large catalogue of mutational signatures that have support from many data sources. At the core of this effort are various tools developed for either *de novo* signature discovery or identifying known mutational signatures in new datasets (refitting). Even though datasets are likely to have a combination of both known and *de novo* signatures, tools that take a combined approach of simultaneous inference of both known and *de novo* signatures have begun to emerge only recently.

We present ReDeNovo, a new method for identification of both known and *de novo* mutational signatures. ReDeNovo demonstrates superior performance over other methods on extensive, carefully-designed synthetic data. Even with increasing levels of noise in the synthetic datasets, our tests show that ReDeNovo is resilient to the inference of false positive signatures as well as to activity estimation errors in patients — an issue that several other tested tools are vulnerable to. Our method additionally demonstrated robustness to variations in parameter settings (Supplementary Section 4). Our tests also revealed that ReDeNovo was the only method that consistently discovered every *de novo* signature. Using ReDeNovo, we have been able to discover a new UV-light related signature. Additionally, ReDeNovo discovers SBS39 in breast and kidney cancers, allowing us to provide additional support for its proposed etiology.

A potential limitation of the current implementation of ReDeNovo is that, theoretically, a *de novo* signature discovered could be a linear combination of known signatures. Although we have yet to encounter such a situation (as ReDeNovo did not infer false positives in our tests on synthetic data), in future we aim to extend ReDeNovo to test for and resolve such ambiguities.

We conclude that the ReDeNovo is a new and powerful tool to uncover mutational signatures, especially in cases where a mixture of known and *de novo* signatures are expected.

## Acknowledgments

Z.N.K., E.H., J.H., B.A., M.G.H., and T.M.P. are supported by the NLM Intramural Research Program. Cancer datasets used in this study were downloaded from the ICGC PCAWG Data Portal (http://dcc.icgc.org/pcawg/). See [26] for detailed descriptions of data and access policies. The results shown here are in whole or part based upon data generated by the TCGA Research Network: https://www.cancer.gov/tcga. This work utilized the computational resources of the NIH HPC Biowulf cluster. (http://hpc.nih.gov)

This research was supported by the Intramural Research Program of the National Institutes of Health (NIH). The contributions of the NIH author(s) are considered Works of the United States Government. The findings and conclusions presented in this paper are those of the author(s) and do not necessarily reflect the views of the NIH or the U.S. Department of Health and Human Services.

## Ethics declaration

The authors declare no competing interests.

