## Supplementary File for "Combined inference of known and novel mutational signatures with ReDeNovo"

### Supplementary Figures

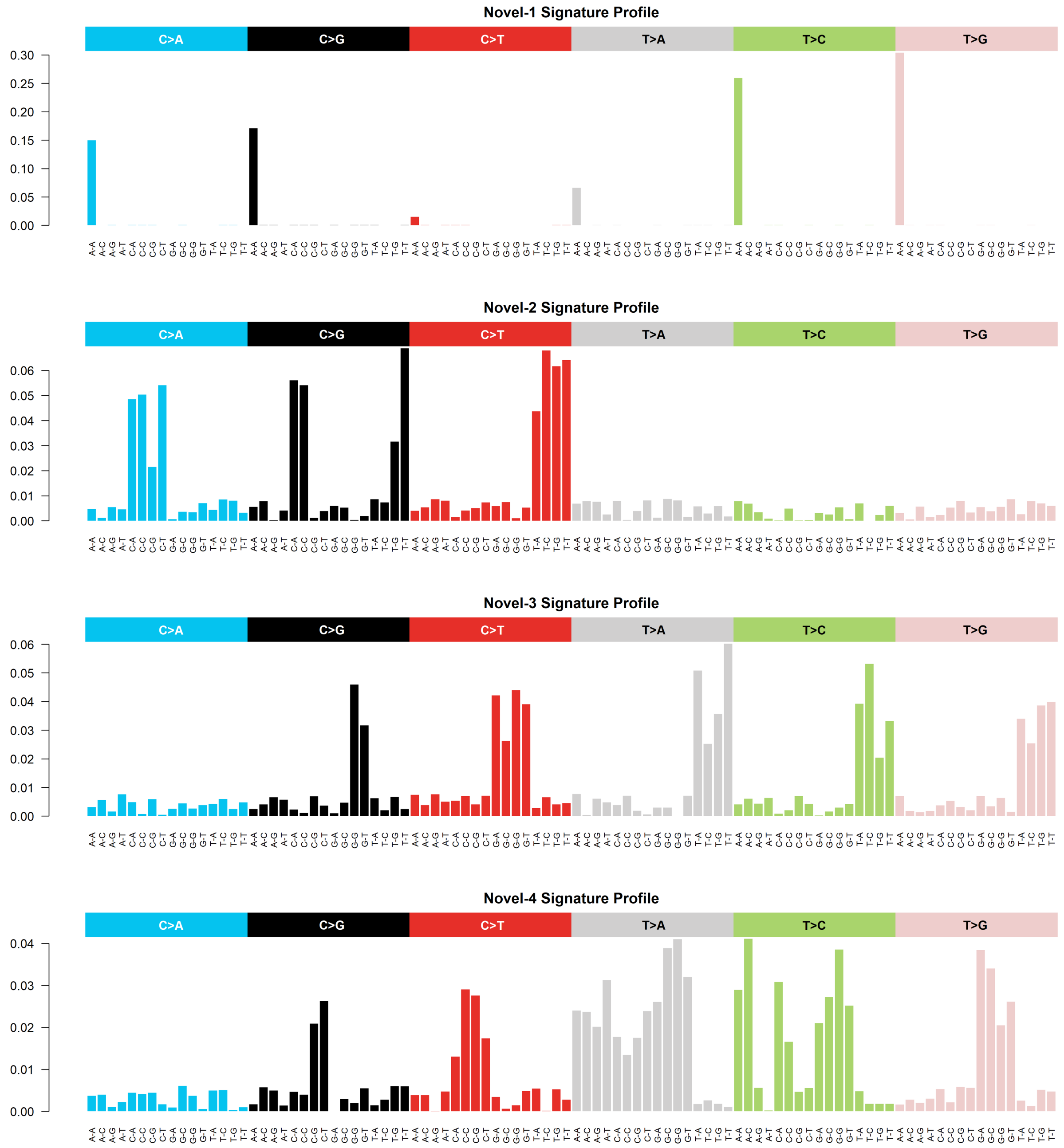


Figure 1: Signature profiles of synthesized signatures.

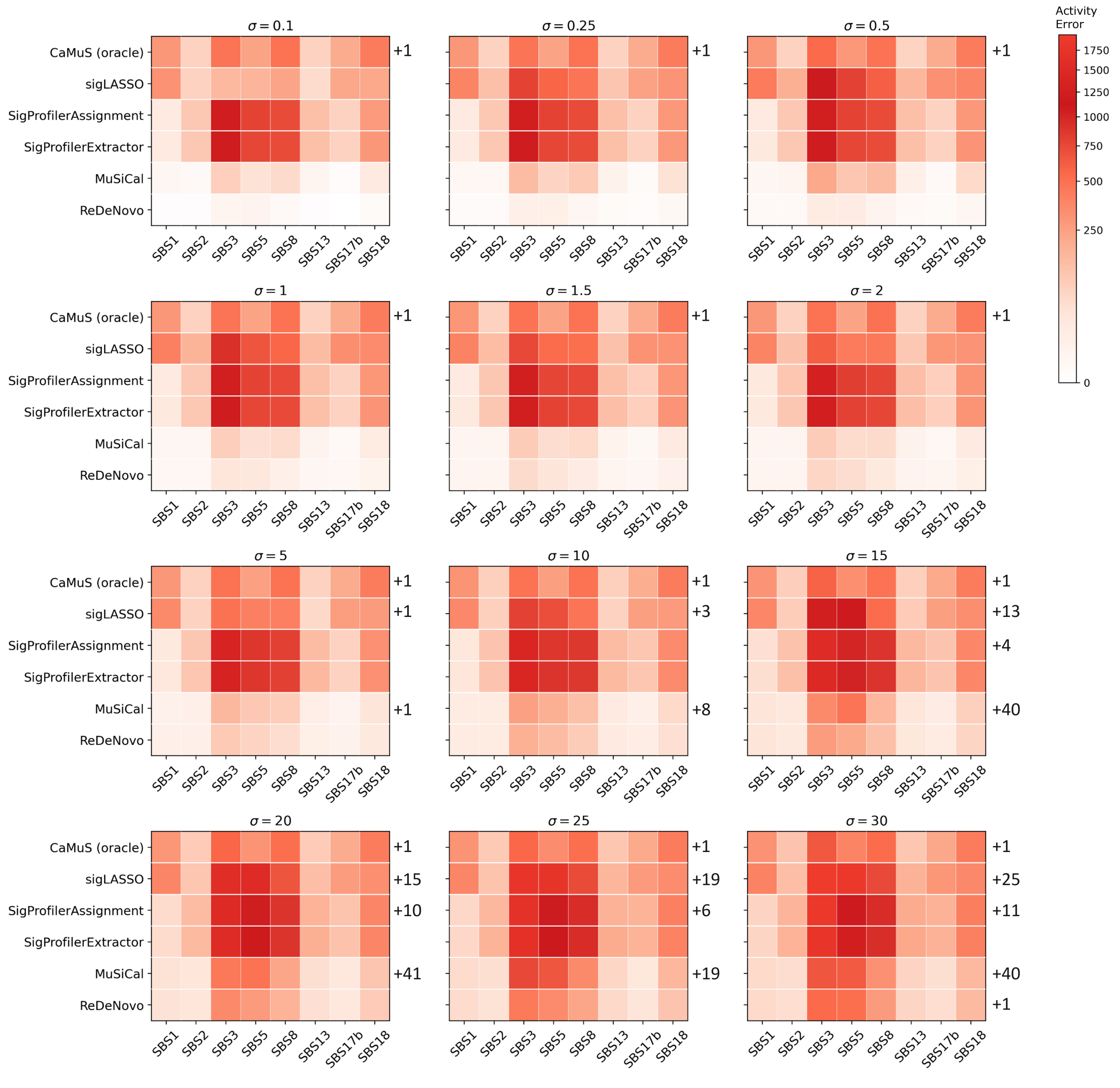


Figure 2: **Performance comparison on noisy synthetic data generated using COSMIC mutational signatures.** Per-signature activity error comparison for noisy synthetic data ($\sigma > 0$) generated using COSMIC mutation signatures. Root mean square error (RMSE) is calculated for activity estimates of each signature (shown individually in each column), averaged over 10 runs to account for inherent stochasticity of the tested tools. The values shown next to each row (e.g., +1, +6) denote the number of additional catalogue signatures that the corresponding method mistakenly inferred in at least one run. These represent false positive signatures that were not present in the ground truth signature set.

#### Supplementary Tables

| Method | $\alpha = 0$ | $\alpha = 0.1$ | $\alpha = 0.25$ | $\alpha = 0.5$ | $\alpha = 1$ | $\alpha = 1.5$ | $\alpha = 2$ |
|---|---|---|---|---|---|---|---|
| CaMuS (oracle) | 0.933±0.000 | 0.933±0.000 | 0.933±0.000 | 0.928±0.000 | 0.933±0.000 | 0.933±0.000 | 0.928±0.000 |
| sigLASSO | **1.000±0.000** | **1.000±0.000** | **1.000±0.000** | **1.000±0.000** | **1.000±0.000** | **1.000±0.000** | **1.000±0.000** |
| SigProfilerAssignment | **1.000±0.000** | **1.000±0.000** | **1.000±0.000** | **1.000±0.000** | **1.000±0.000** | **1.000±0.000** | **1.000±0.000** |
| SigProfilerExtractor | **1.000±0.000** | **1.000±0.000** | **1.000±0.000** | **1.000±0.000** | **1.000±0.000** | **1.000±0.000** | **1.000±0.000** |
| MuSiCal | **1.000±0.000** | **1.000±0.000** | **1.000±0.000** | **1.000±0.000** | **1.000±0.000** | **1.000±0.000** | **1.000±0.000** |
| ReDeNovo | **1.000±0.000** | **1.000±0.000** | **1.000±0.000** | **1.000±0.000** | **1.000±0.000** | **1.000±0.000** | **1.000±0.000** |

| Method | $\alpha = 5$ | $\alpha = 10$ | $\alpha = 15$ | $\alpha = 20$ | $\alpha = 25$ | $\alpha = 30$ |
|---|---|---|---|---|---|---|
| CaMuS (oracle) | 0.933±0.000 | 0.933±0.000 | 0.928±0.000 | 0.933±0.000 | 0.875±0.000 | 0.933±0.000 |
| sigLASSO | 0.941±0.000 | 0.842±0.000 | 0.576±0.012 | 0.518±0.005 | 0.479±0.010 | 0.413±0.009 |
| SigProfilerAssignment | **1.000±0.000** | **1.000±0.000** | 0.800±0.000 | 0.615±0.000 | 0.727±0.000 | 0.593±0.000 |
| SigProfilerExtractor | **1.000±0.000** | **1.000±0.000** | **1.000±0.000** | **1.000±0.000** | **1.000±0.000** | **1.000±0.000** |
| MuSiCal | 0.962±0.115 | 0.988±0.024 | 0.865±0.175 | 0.800±0.179 | 0.725±0.186 | 0.862±0.137 |
| ReDeNovo | **1.000±0.000** | **1.000±0.000** | **1.000±0.000** | **1.000±0.000** | **1.000±0.000** | 0.941±0.000 |

Table 1: **Performance comparison for recovering the ground truth signature set on synthetic data generated using COSMIC mutational signatures across different noise levels.** Performance is measured using the F1 score for recovery of the ground-truth signature set. Each column represents a different level of standard deviation of Gaussian noise in the input mutation count data. F1 score is averaged over 10 runs to account for inherent stochasticity of the tested tools.

| Method | $\alpha = 0$ | $\alpha = 0.1$ | $\alpha = 0.25$ | $\alpha = 0.5$ | $\alpha = 1$ | $\alpha = 1.5$ | $\alpha = 2$ |
|---|---|---|---|---|---|---|---|
| CaMuS (oracle) | 98.8±0.0 | 98.8±0.0 | 98.8±0.0 | 112.1±39.9 | 98.7±0.0 | 99.0±0.0 | 112.5±39.7 |
| sigLASSO | 55.4±0.0 | 60.2±1.0 | 131.7±2.4 | 176.3±1.8 | 155.3±0.7 | 131.4±1.1 | 114.0±0.8 |
| SigProfilerAssignment | 178.0±0.0 | 179.3±0.0 | 180.1±0.0 | 180.3±0.0 | 181.4±0.0 | 181.3±0.0 | 189.0±0.0 |
| SigProfilerExtractor | 177.8±0.0 | 177.8±0.0 | 176.9±0.0 | 178.3±0.0 | 178.5±0.0 | 181.6±0.0 | 184.9±0.0 |
| MuSiCal | 16.8.±13.9 | 15.9±14.9 | 13.1±11.2 | 19.1±13.5 | 36.1±32.0 | 24.1±36.2 | 14.5±10.1 |
| ReDeNovo | **0.2±0.0** | **0.6±0.1** | **1.5±0.2** | **2.4±0.1** | **4.7±0.1** | **7.0±0.2** | **9.1±0.2** |

| Method | $\alpha = 5$ | $\alpha = 10$ | $\alpha = 15$ | $\alpha = 20$ | $\alpha = 25$ | $\alpha = 30$ |
|---|---|---|---|---|---|---|
| CaMuS (oracle) | 99.7±0.0 | 100.3±0.0 | 116.6±40.9 | 107.7±0.0 | 109.2±0.0 | 117.3±0.0 |
| sigLASSO | 100.1±1.0 | 137.8±0.4 | 203.3±1.0 | 262.2±1.5 | 300.7±0.5 | 323.4±0.8 |
| SigProfilerAssignment | 199.1±0.0 | 207.4±0.0 | 225.0±0.0 | 228.8±0.0 | 249.4±0.0 | 272.2±0.0 |
| SigProfilerExtractor | 198.6±0.0 | 208.5±0.0 | 223.7±0.0 | 233.8±0.0 | 247.5±0.0 | 263.9±0.0 |
| MuSiCal | 27.1±8.2 | 34.2±8.9 | 53.6±23.8 | 88.1±36.3 | 150.1±53.5 | 108.1±23.9 |
| ReDeNovo | **14.0±0.0** | **27.3±0.0** | **40.1±0.0** | **53.6±0.0** | **67.7±0.0** | **100.9±0.0** |

Table 2: **Performance comparison for activity errors on synthetic data generated using COSMIC mutational signatures across different noise levels.** Performance is measured using the RMSE of all the inferred activities. Each column represents a different level of standard deviation of Gaussian noise in the input mutation count data. F1 scores are averaged over 10 runs to account for inherent stochasticity of the tested tools.

Table 3: **Performance comparison on additional 15 synthetic datasets generated using COSMIC mutational signatures.** Table includes F1 scores for recovering the ground truth signature set and RMSE of all the inferred activities. Both reported measures are averaged over 10 runs to account for inherent stochasticity of the tested tools. [RMSE: root mean square error]

| **Simulated Data** | **Method** | **RMSE** | **F1 Score** |
|---|---|---|---|
| Synthetic dataset 2 | CaMuS | 497.6±0.0 | 0.8000±0.0000 |
| | sigLASSO | 110.0±0.0 | **1.0000±0.0000** |
| | SigProfilerAssignment | 244.4±0.0 | 0.8889±0.0000 |
| | SigProfilerExtractor | 253.8±0.0 | **1.0000±0.0000** |
| | MuSiCal | 261.8±184.6 | **1.0000±0.0000** |
| | ReDeNovo | **0.2±0.0** | **1.0000±0.0000** |
| Synthetic dataset 3 | CaMuS | 486.6±0.0 | 0.6250±0.0000 |
| | sigLASSO | 28.0±0.0 | **1.0000±0.0000** |
| | SigProfilerAssignment | 86.2±0.0 | **1.0000±0.0000** |
| | SigProfilerExtractor | 102.3±0.0 | **1.0000±0.0000** |
| | MuSiCal | 252.9±233.9 | **1.0000±0.0000** |
| | ReDeNovo | **0.2±0.0** | **1.0000±0.0000** |
| Synthetic dataset 4 | CaMuS | 603.4±0.0 | 0.7826±0.0000 |
| | sigLASSO | 91.4±0.0 | **1.0000±0.0000** |
| | SigProfilerAssignment | 307.9±0.0 | **1.0000±0.0000** |
| | SigProfilerExtractor | 315.3±0.0 | **1.0000±0.0000** |
| | MuSiCal | 100.8±23.7 | **1.0000±0.0000** |
| | ReDeNovo | **0.2±0.0** | **1.0000±0.0000** |
| Synthetic dataset 5 | CaMuS | 595.8±0.1 | 0.7273±0.0000 |
| | sigLASSO | 22.4±0.0 | **1.0000±0.0000** |
| | SigProfilerAssignment | 16.2±0.0 | **1.0000±0.0000** |
| | SigProfilerExtractor | 23.2±0.0 | **1.0000±0.0000** |
| | MuSiCal | 69.1±97.0 | **1.0000±0.0000** |
| | ReDeNovo | **0.1±0.0** | **1.0000±0.0000** |
| Synthetic dataset 6 | CaMuS | 267.2±0.0 | 0.8571±0.0000 |
| | sigLASSO | 18.3±0.0 | **1.0000±0.0000** |
| | SigProfilerAssignment | 34.9±0.0 | **1.0000±0.0000** |
| | SigProfilerExtractor | 40.1±0.0 | **1.0000±0.0000** |
| | MuSiCal | 19.1±9.6 | **1.0000±0.0000** |
| | ReDeNovo | **0.2±0.0** | **1.0000±0.0000** |
| Synthetic dataset 7 | CaMuS | 436.2±1.1 | 0.7778±0.0000 |
| | sigLASSO | 50.0±0.0 | **1.0000±0.0000** |
| | SigProfilerAssignment | 93.9±0.0 | **1.0000±0.0000** |
| | SigProfilerExtractor | 203.9±0.0 | **1.0000±0.0000** |
| | MuSiCal | 106.8±199.6 | **1.0000±0.0000** |
| | ReDeNovo | **0.2±0.0** | **1.0000±0.0000** |
| Synthetic dataset 8 | CaMuS | 300.7±0.0 | 0.8000±0.0000 |
| | sigLASSO | 48.1±0.0 | **1.0000±0.0000** |
| | SigProfilerAssignment | 19.9±0.0 | **1.0000±0.0000** |
| | SigProfilerExtractor | 60.7±0.0 | **1.0000±0.0000** |
| | MuSiCal | 61.9±47.1 | **1.0000±0.0000** |
| | ReDeNovo | **0.2±0.0** | **1.0000±0.0000** |
| Synthetic dataset 9 | CaMuS | 332.2±6.9 | 0.9000±0.0000 |
| | sigLASSO | 57.6±0.0 | **1.0000±0.0000** |
| | SigProfilerAssignment | 191.4±0.0 | **1.0000±0.0000** |
| | SigProfilerExtractor | 198.8±0.0 | **1.0000±0.0000** |
| | MuSiCal | 264.3±166.6 | **1.0000±0.0000** |
| | ReDeNovo | **0.2±0.0** | **1.0000±0.0000** |

*Continued on next page*

| Simulated Data | Method | RMSE | F1 Score |
|---|---|---|---|
| Synthetic dataset 10 | CaMuS | 269.6±33.3 | 0.8333±0.0192 |
| | sigLASSO | 28.9±0.0 | **1.0000±0.0000** |
| | SigProfilerAssignment | 89.6±0.0 | **1.0000±0.0000** |
| | SigProfilerExtractor | 117.7±0.0 | **1.0000±0.0000** |
| | MuSiCal | 14.0±15.1 | **1.0000±0.0000** |
| | ReDeNovo | **0.2±0.0** | **1.0000±0.0000** |
| Synthetic dataset 11 | CaMuS | 406.7±20.1 | 0.8333±0.0100 |
| | sigLASSO | 98.8±0.0 | **1.0000±0.0000** |
| | SigProfilerAssignment | 269.2±0.0 | **1.0000±0.0000** |
| | SigProfilerExtractor | 272.0±0.0 | **1.0000±0.0000** |
| | MuSiCal | 226.1±154.0 | 0.9338±0.0415 |
| | ReDeNovo | **0.2±0.0** | **1.0000±0.0000** |
| Synthetic dataset 12 | CaMuS | 532.8±0.8 | 0.7619±0.0000 |
| | sigLASSO | 88.2±0.0 | **1.0000±0.0000** |
| | SigProfilerAssignment | 293.2±0.0 | 0.9524±0.0000 |
| | SigProfilerExtractor | 303.8±0.0 | **1.0000±0.0000** |
| | MuSiCal | 48.8±26.3 | **1.0000±0.0000** |
| | ReDeNovo | **0.2±0.0** | **1.0000±0.0000** |
| Synthetic dataset 13 | CaMuS | 683.9±0.0 | 0.7500±0.0000 |
| | sigLASSO | 119.1±0.1 | **1.0000±0.0000** |
| | SigProfilerAssignment | 254.1±0.0 | 0.9600±0.0000 |
| | SigProfilerExtractor | 264.4±0.0 | **1.0000±0.0000** |
| | MuSiCal | 200.4±90.4 | **1.0000±0.0000** |
| | ReDeNovo | **0.2±0.0** | **1.0000±0.0000** |
| Synthetic dataset 14 | CaMuS | 525.3±0.0 | 0.7000±0.0000 |
| | sigLASSO | 82.4±0.0 | **1.0000±0.0000** |
| | SigProfilerAssignment | 234.5±0.0 | **1.0000±0.0000** |
| | SigProfilerExtractor | 239.8±0.0 | **1.0000±0.0000** |
| | MuSiCal | 148.9±50.6 | **1.0000±0.0000** |
| | ReDeNovo | **0.2±0.0** | **1.0000±0.0000** |
| Synthetic dataset 15 | CaMuS | 458.8±17.4 | 0.8800±0.0262 |
| | sigLASSO | 117.3±0.1 | **1.0000±0.0000** |
| | SigProfilerAssignment | 74.2±0.0 | **1.0000±0.0000** |
| | SigProfilerExtractor | 82.2±0.0 | **1.0000±0.0000** |
| | MuSiCal | 183.7±109.3 | **1.0000±0.0000** |
| | ReDeNovo | **0.2±0.0** | **1.0000±0.0000** |
| Synthetic dataset 16 | CaMuS | 460.4±0.0 | 0.7826±0.0000 |
| | sigLASSO | 88.3±0.0 | **1.0000±0.0000** |
| | SigProfilerAssignment | 224.7±0.0 | **1.0000±0.0000** |
| | SigProfilerExtractor | 231.1±0.0 | **1.0000±0.0000** |
| | MuSiCal | 251.0±0.0 | **1.0000±0.0000** |
| | ReDeNovo | **0.2±0.0** | **1.0000±0.0000** |

| | Method | Synth-1 | Synth-2 | Synth-3 | Synth-4 | Hidden SBS1 | Hidden SBS2 | Hidden SBS3 | Hidden SBS5 | Hidden SBS8 | Hidden SBS13 | Hidden SBS17b | Hidden SBS18 | Average |
|---|---|---|---|---|---|---|---|---|---|---|---|---|---|---|
| **Detect of *de novo* sig** | CaMuS (oracle) | 1.00 ✓ | 1.00 ✓ | 1.00 ✓ | 1.00✓ | ✗ | 1.00 ✓ | ✗ | 1.00 ✓ | 1.00 ✓ | 1.00 ✓ | 0.90✓ | ✗ | 0.74 |
| | SigProfilerExtractor | 1.00 ✓ | ✗ | 1.00 ✓ | 1.00✓ | | | | | | | | | 0.75 |
| | MuSiCal | 0.10 ✗ | ✗ | ✗ | ✗ | ✗ | ✗ | ✗ | ✗ | ✗ | 1.00 ✓ | 1.00 ✓ | ✗ | 0.18 |
| | ReDeNovo | 1.00 ✓ | 1.00 ✓ | 1.00 ✓ | 1.00 ✓ | 1.00 ✓ | 1.00 ✓ | 1.00 ✓ | 1.00 ✓ | 1.00 ✓ | 1.00 ✓ | 1.00 ✓ | 1.00 ✓ | **1.00** |
| **Cos sim of *de novo* sig** | CaMuS (oracle) | 0.98±0.00 | 0.89±0.00 | 0.92±0.00 | 0.95±0.00 | ✗ | 0.98±0.00 | ✗ | 0.92±0.00 | 0.91±0.00 | 0.99±0.00 | 0.99±0.30 | ✗ | 0.703 |
| | SigProfilerExtractor | 1.00±0.00 | ✗ | 1.00±0.00 | 1.00±0.00 | | | | | | | | | 0.750 |
| | MuSiCal | 0.10±0.30 | ✗ | ✗ | ✗ | ✗ | ✗ | ✗ | ✗ | ✗ | 0.99±0.00 | 0.97±0.01 | ✗ | 0.164 |
| | ReDeNovo | 0.99±0.00 | 0.93±0.00 | 0.97±0.00 | 0.98±0.00 | 0.83±0.03 | 0.94±0.02 | 0.91±0.01 | 0.88±0.00 | 0.92±0.00 | 0.87±0.02 | 0.98±0.00 | 0.87±0.00 | **0.918** |
| **F1 score all sigs** | CaMuS (oracle) | 0.737±0.000 | 0.800±0.000 | 0.706±0.000 | 0.706±0.000 | 0.787±0.040 | 0.625±0.000 | 0.857±0.000 | **0.933±0.000** | 0.933±0.000 | 0.375±0.000 | 0.821±0.007 | 0.933±0.000 | 0.768 |
| | SigProfilerExtractor | **1.000±0.000** | 0.762±0.000 | **1.000±0.000** | **1.000±0.000** | | | | | | | | | 0.941 |
| | MuSiCal | 0.947±0.018 | 0.802±0.107 | 0.868±0.094 | 0.657±0.100 | 0.870±0.015 | 0.746±0.073 | 0.704±0.134 | 0.588±0.085 | 0.804±0.056 | **0.900±0.115** | **1.000±0.000** | 0.854±0.025 | 0.811 |
| | ReDeNovo | 0.941±0.000 | **0.988±0.024** | **1.000±0.000** | 0.941±0.000 | **1.000±0.000** | **0.773±0.048** | **1.000±0.000** | **0.933±0.000** | **1.000±0.000** | 0.769±0.000 | 0.985±0.031 | **1.000±0.000** | **0.944** |
| **Act. RMSE all sigs** | CaMuS (oracle) | 251.3±0.3 | 180.0±0.0 | 278.5±0.0 | 215.3±0.0 | 352.9±8.3 | 510.7±0.1 | 399.0±0.4 | 160.5±0.0 | 173.1±0.0 | 584.3±0.2 | 309.4±17.0 | 249.3±0.0 | 305.4 |
| | SigProfilerExtractor | 171.1±1.7 | 472.2±0.0 | 195.4±4.0 | 205.7±2.0 | | | | | | | | | 261.1 |
| | MuSiCal | 283.8±63.2 | 197.1±48.0 | 259.5±14.0 | 264.2±60.7 | 139.3±32.5 | 795.1±15.8 | 379.1±7.0 | 261.4±9.4 | 151.8±7.7 | 450.9±417.9 | 66.7±8.7 | 66.8±23.0 | 277.9 |
| | ReDeNovo | **119.4±0.0** | **56.6±10.1** | **39.0±0.0** | **81.1±0.0** | **111.4±8.5** | **386.7±55.9** | **109.1±5.0** | **104.0±1.3** | **61.2±1.8** | **415.9±23.3** | **28.1±24.6** | **52.7±0.0** | **130.4** |

Table 4: **Performance comparison on synthetic data with COSMIC signatures and one *de novo* signature.** Each column shows results for a different *de novo* signature in the data, combined with the same fixed set of COSMIC signatures (SBS1, SBS2, SBS3, SBS5, SBS8, SBS13, SBS17b, SBS18). The first four *de novo* signatures are synthetically generated. To make the test as realistic as possible, we also individually hid each COSMIC signature from the input catalogue and treated it as a *de novo* signature. We treat the predicted *do novo* signature as accurately detected if its signature profile has a cosine similarity of at least 0.70 with the ground truth *de novo* signature profile. This table reports the detection rate of *de novo* signatures, the cosine similarity between real and inferred *de novo* signatures, the F1 score for the presence/absence of all signatures named "F1 score all sigs," and the RMSE of all the inferred activities. All reported measures are averaged (mean) over 10 runs to account for inherent stochasticity of the tested tools. Symbol ✗ indicates cases where the signature was not inferred; gray shading denotes not applicable.

Table 5: Key literature references, with comments.

| Cancer type | Signature | Sample of references |
|---|---|---|
| Breast-AdenoCA | SBS1 | [1], [2], [3] |
| | SBS2 | [1], [2], [3] |
| | SBS3 | [1], [2], [3] |
| | SBS5 | [1], [2], [3] |
| | SBS8 | [1], [2], [3] |
| | SBS9 | [1], [4], [5] |
| | SBS13 | [1], [2], [3] |
| | SBS18 | [1], [2], [3] |
| | SBS39 | Sig. not found, relation to HRD [6], relation to NHEJ deficiency [7] |
| | SBS40a/c | [1], [8] |
| CNS-Medullo | SBS1 | [1],[9] |
| | SBS3 | [9] |
| | SBS5 | [1], [9] |
| | SBS8 | [1] |
| | SBS15 | Sig not found; presence of MMRd [10] |
| | SBS18 | [1], [9] |
| | SBS31 | [1] [11] |
| | SBS39 | [1] |
| | SBS40a/c | [1], [9] |
| | SBS46 | Sig not found; sequencing artifact |
| Kidney-RCC | SBS1 | [1], [12], [13] |
| | SBS3 | [14] |
| | SBS4 | [15], [16], [13] |
| | SBS12 | [16] |
| | SBS18 | [16], [13] |
| | SBS39 | Not found; not found, relation to HRD [6], relation to NHEJ deficiency [7] |
| | SBS40a/b/c | [1], [12], [16], [13] |
| Liver-HCC | SBS3 | [17], [18] |
| | SBS5 | [1], [15], [18] |
| | SBS6 | [1], [15] |
| | SBS8 | [19] |
| | SBS12 | [1], [15], [18] |
| | SBS16 | [1] [15] |
| | SBS25 | [18] |
| | SBS40a/b | [1], [18], [17] |
| | SBS92 | [14] |
| | SBS95 | Sequencing artifact |
| Lymph-BNHL | SBS1 | [1], [20], [21] |
| | SBS3 | [1], [20], [21] |
| | SBS5 | [1], [22], [21] |
| | SBS6 | [1] |
| | SBS9 | [1], [20], [22], [21] |
| | SBS10c | Sig. not found |
| | SBS17b | [1], [20], [22], [21], [23] |

*Continued on next page*

| Cancer type | Signature | Sample of Reference |
|---|---|---|
| | SBS25 | Sig. not found Non Hodgkin's lymphoma but related to chemotherapy in classic Hodgkin's lymphoma [1][24] |
| | SBS40a | [1] |
| | SBS46 | Sig. not found; sequencing artifact |
| | SBS85 | [25], [26] |
| | SBS93 | Sig. not found; etiology unknown |
| Ovary-AdenoCA | SBS1 | [1], [27], [28], [29] |
| | SBS3 | [1], [27], [28], [29] |
| | SBS5 | [1], [27], [28], [29] |
| | SBS8 | [1], [29] |
| | SBS13 | [1], [28], [29] |
| | SBS37 | [29] |
| | SBS39 | [1], [29] |
| | SBS40a/c | [1], [27] |
| | SBS89 | Sig. not found, etiology unknown |
| Panc-AdenoCA | SBS1 | [1], [30], [31], [32], [33] |
| | SBS2 | [1], [31] |
| | SBS3 | [34], [30], [31], [35], [36], [32], [33] |
| | SBS5 | [1], [32] |
| | SBS8 | [1], [30] |
| | SBS9 | [37] |
| | SBS10c | [38] |
| | SBS13 | [1], [31] |
| | SBS15 | Sig. not found, evidence of MMRd [39] |
| | SBS40a | [1] |
| Prost-AdenoCA | SBS1 | [1], [40] |
| | SBS3 | [1], [40] |
| | SBS5 | [1], [40] |
| | SBS6 | [40], [41] |
| | SBS8 | [1] |
| | SBS37 | [1] |
| | SBS40a/c | [1] |
| | SBS41 | [1] |
| Skin-Melanoma | SBS2 | [1], [42] |
| | SBS7a/b | [1], [17], [42], [43] |

Table 6: **Hyperparameter tuning on synthetic data** Table includes F1 score for recovering the ground truth signature set and RMSE of all the inferred activities, both averaged over 10 runs. [RMSE: root mean square error]

| Parameter | Value | RMSE | F1 Score |
|---|---|---|---|
| consno (default = 1) | 2 | 0.2±0.0 | 1.0000±0.0000 |
| | 3 | 0.2±0.0 | 1.0000±0.0000 |
| | 4 | 0.2±0.0 | 1.0000±0.0000 |
| | 5 | 79.2±8.9 | 0.8952±0.0381 |
| numiters (default = 10) | 2 | 0.2±0.0 | 1.0000±0.0000 |
| | 3 | 0.2±0.0 | 1.0000±0.0000 |
| | 4 | 0.2±0.0 | 1.0000±0.0000 |
| | 5 | 0.2±0.0 | 1.0000±0.0000 |
| | 6 | 0.2±0.0 | 1.0000±0.0000 |
| | 7 | 0.2±0.0 | 1.0000±0.0000 |
| | 8 | 0.2±0.0 | 1.0000±0.0000 |
| | 9 | 0.2±0.0 | 1.0000±0.0000 |
| numruns (default = 10) | 2 | 0.2±0.0 | 1.0000±0.0000 |
| | 3 | 0.2±0.0 | 1.0000±0.0000 |
| | 4 | 0.2±0.0 | 1.0000±0.0000 |
| | 5 | 0.2±0.0 | 1.0000±0.0000 |
| | 6 | 0.2±0.0 | 1.0000±0.0000 |
| | 7 | 0.2±0.0 | 1.0000±0.0000 |
| | 8 | 0.2±0.0 | 1.0000±0.0000 |
| | 9 | 0.2±0.0 | 1.0000±0.0000 |
| thr1 (default = 0.10) | 0.01 | 0.2±0.0 | 1.0000±0.0000 |
| | 0.02 | 0.2±0.0 | 1.0000±0.0000 |
| | 0.05 | 0.2±0.0 | 1.0000±0.0000 |
| | 0.15 | 70.3±0.0 | 0.9333±0.0000 |
| | 0.20 | 88.1±0.0 | 0.8571±0.0000 |
| thr2 (default = 0.70) | 0.75 | 0.2±0.0 | 1.0000±0.0000 |
| | 0.80 | 0.2±0.0 | 1.0000±0.0000 |
| | 0.85 | 0.2±0.0 | 1.0000±0.0000 |
| thr3 (default = 0.70) | 0.60 | 0.2±0.0 | 1.0000±0.0000 |
| | 0.80 | 0.2±0.0 | 1.0000±0.0000 |
| | 0.90 | 0.2±0.0 | 1.0000±0.0000 |
| | 1.00 | 0.2±0.0 | 1.0000±0.0000 |
| thr4 (default = (0.80) | 0.70 | 0.2±0.0 | 1.0000±0.0000 |
| | 0.75 | 0.2±0.0 | 1.0000±0.0000 |
| | 0.85 | 0.2±0.0 | 1.0000±0.0000 |
| | 0.90 | 0.2±0.0 | 1.0000±0.0000 |
| thr5 (default = 0.10) | 0.01 | 0.2±0.0 | 1.0000±0.0000 |
| | 0.02 | 0.2±0.0 | 1.0000±0.0000 |
| | 0.05 | 0.2±0.0 | 1.0000±0.0000 |
| | 0.15 | 0.2±0.0 | 1.0000±0.0000 |
| output-exposure-thr1 (default = 0.05) | 0.01 | 0.2±0.0 | 1.0000±0.0000 |
| | 0.02 | 0.2±0.0 | 1.0000±0.0000 |
| | 0.15 | 447.0±0.0 | 0.5455±0.0000 |
| | 0.20 | 447.0±0.0 | 0.5455±0.0000 |
| | 0.25 | 447.0±0.0 | 0.5455±0.0000 |
| output-exposure-thr2 (default = 1) | 5 | 0.2±0.0 | 1.0000±0.0000 |
| | 10 | 0.2±0.0 | 1.0000±0.0000 |
| | 25 | 0.2±0.0 | 1.0000±0.0000 |
| | 50 | 0.2±0.0 | 1.0000±0.0000 |
| | 100 | 0.2±0.0 | 1.0000±0.0000 |

### Supplementary Materials

### 1 Simulation Data Generation

#### 1.1 Simulated dataset with varying levels of noise

Simulated data was generated to test if the ReDeNovo can uncover novel signatures and their activities in addition to the known signatures. The mutation count matrix $M$ that serves as an input to our method was generated as specified by the NMF formulation in the Methods section of the main manuscript. In order to compute $M_{n\times m}$, the two low rank matrices $A_{n\times l}$ and $P_{l\times m}$ were simulated as described below. These matrices are then used as the ground truth to test the performance of ReDeNovo and other methods. To evaluate the robustness of the methods, we added varying levels of noise $R_{n\times m}$ to $M$. In this simulation, we assumed that $n = 198$ (number of patients in the real dataset) and $m = 96$ (number of mutation categories).

**Signature matrix** $P_{n\times m}$: To construct the signature matrix $P$, we used COSMIC v.3.4 signatures for the existing signatures.

**Activity matrix** $A_{n\times l}$: We generated 16 synthetic cancer datasets with distinct sets of signatures (listed below). For each dataset, activity score matrix $A$ was generated from a multivariate truncated normal distribution with different mean $\mu$ and covariance matrix $\Sigma$ for each dataset (the resulting numbers scaled to range from 0 to 10,000).

Synthetic dataset 1 was generated using the same signatures set as detected by ReDeNovo in Breast cancer data. For the other 15, we used signatures used by MuSiCal [44] for their synthetic experiments. For the synthetic dataset 1, we are able to use the estimated $\mu$ and $\Sigma$ values. For the remaining 15 datasets, we do not have mutation counts and activity scores for the chosen signatures, so we draw $\mu$ and $\Sigma$ uniformly at random from value intervals observed in synthetic dataset 1. Since such randomly generated covariance matrices are not necessarily positive semi-definite, we applied methods from [45] to obtain a positive semi-definite approximation of the randomly generated covariance matrices.

**Activity matrix sparsification:** For each signature, we randomly select 10% to 30% of the samples and set their activity scores to 0.

**Noise matrix** $R$: The mutation count noise matrix $R_{n\times m}$ is generated from a normal distribution with mean 0 and standard deviation $\sigma = 0.1, 0.25, 0.5, 1.0, 1.5, 2.0, 5.0, 10.0, 15.0, 20.0, 25.0$, and $30.0$. Then, the mutation count matrix for each synthetic dataset is calculated as $M = A \times P + R$. In case that adding the $R$ matrix results in negative mutation counts, we set those negative values to 0.

***De novo* signature**: We generated 4 synthetic signature profiles for the COSMIC v.3.4 mutational categories and the corresponding activities. For each *de novo* signature, its mutational profile is drawn from a uniform distribution between 0 to 0.1 as well as manually assigned higher values representing more frequent mutation categories in the signature profile. Only for synthetic signature 1, we used manually defined peaks. Finally, the signature profile is normalized to sum up to 1 (See Supplementary Figure 1). The activity score matrix of the 4 novel signatures is generated using the same simulation schema.

**Signatures used in the 16 datasets:**

Synthetic dataset 1 signatures: [SBS1, SBS2, SBS3, SBS5, SBS8, SBS13, SBS17b, SBS18]

The other 15 simulated datasets were generated from cancer types using common signature sets from MuSiCal [44], adapted to COSMIC version 3.4. The signature sets are as follows:

Synthetic dataset 2 signatures: [SBS1, SBS2, SBS3, SBS5, SBS12, SBS13, SBS17a, SBS17b, SBS18, SBS22a, SBS24, SBS32]

Synthetic dataset 3 signatures: [SBS1, SBS2, SBS5, SBS8, SBS13, SBS29]

Synthetic dataset 4 signatures: [SBS1, SBS2, SBS3, SBS5, SBS8, SBS13, SBS17a, SBS17b, SBS18, SBS37, SBS41]

Synthetic dataset 5 signatures: [SBS1, SBS5, SBS11, SBS30, SBS37]

Synthetic dataset 6 signatures: [SBS1, SBS5, SBS8, SBS18, SBS39, SBS60]

Synthetic dataset 7 signatures: [SBS1, SBS5, SBS17a, SBS17b, SBS18, SBS28, SBS37, SBS45]

Synthetic dataset 8 signatures: [SBS1, SBS2, SBS5, SBS13, SBS22a, SBS29, SBS41]

Synthetic dataset 9 signatures: [SBS1, SBS2, SBS3, SBS4, SBS5, SBS13, SBS17a, SBS17b, SBS18]

Synthetic dataset 10 signatures: [SBS1, SBS2, SBS4, SBS5, SBS8, SBS13]

Synthetic dataset 11 signatures: [SBS1, SBS2, SBS3, SBS5, SBS13, SBS17a, SBS17b, SBS34, SBS36, SBS37, SBS56]

Synthetic dataset 12 signatures: [SBS1, SBS2, SBS3, SBS5, SBS8, SBS13, SBS18, SBS35, SBS39, SBS41]

Synthetic dataset 13 signatures: [SBS1, SBS2, SBS3, SBS5, SBS8, SBS13, SBS17a, SBS17b, SBS18, SBS28, SBS30, SBS51]

Synthetic dataset 14 signatures: [SBS1, SBS2, SBS3, SBS5, SBS13, SBS18, SBS33, SBS37, SBS41, SBS52]

Synthetic dataset 15 signatures: [SBS1, SBS2, SBS5, SBS7a, SBS7b, SBS7c, SBS7d, SBS13, SBS17a, SBS17b, SBS38, SBS58]

Synthetic dataset 16 signatures: [SBS1, SBS2, SBS3, SBS5, SBS13, SBS17a, SBS17b, SBS18, SBS41, SBS51, SBS58]

### 2 Gradient Descent Details

This section describes the gradient descent optimization subroutine leveraged by our ReDeNovo pipeline. This logic is implemented in Python 3.10 using the TensorFlow Keras API ≥ v2.15 [46]. We compute the Frobenius norm between $M$ and $A \times P$ as loss function, where $P$ constitutes a mixture of fixed signatures and inferred signatures. By default, we employ the adaptive moment estimation method (Adam, [47]) in combination with a custom learning rate strategy to fit our model to the given data. We found Adam to provide an appropriate balance between performance and accuracy in the context of sparse data, however, our method supports any additional estimators available via the Keras API.

Starting with random values for $A$ and the components to be inferred in $P$, the gradients for these matrices are computed according to the loss function and scaled based on the user-specified learning rate as provided by Adam. These gradients are consequently applied to the corresponding matrices and the process is repeated a user defined number of times. To reduce the probability of converging to a local minima during the gradient descent optimization, we bracket our iterations into a series of batches defined by varying learning rate and number of iterations where each consecutive batch starts from the best solution from the current batch according to the loss. Optionally, a different optimizer can be chosen for any such bracket. This approach allows for estimating suitable values for $A$ and $P$ by starting with initially larger learning rates that can be gradually reduced as the optimization begins to converge. While tuning the learning rate and number of iterations may be necessary based on the characteristics of the input data, we provide a default strategy used throughout the manuscript that has shown to be reliable and effective in producing consistent, interpretable outcomes.

Note that mutational signature inference corresponds to a constrained optimization problem in which signatures in $P$ must conform to probability distributions. To account for this in Tensorflow, we apply a projection at each iteration that clips negative values to 0 and normalizes each signature in $P$ by the sum of its values.Together with the optimization strategy described above, this allows ReDeNovo to harness TensorFlow's optimization capabilities while maintaining convergence to biologically valid and meaningful solution.

In our experiments, we employ nine epochs, each with 5,000 steps. The step sizes of each epoch are 10, 1, 0.1, 0.1, 0.01, 0.01, 0.01, 0.001, 0.0001.

Also see Supplementary Materials Section 4 for other default parameters of ReDeNovo.

### 3 ReDeNovo Algorithm

---

**Algorithm 1** ReDeNovo

---

**Input:** nRuns, nIters, initSigs, refSigs $M$

novelSigs ← []
runSigs ← []
**for** run in nRuns **do**
  knownSigs ← initSig
  banned ← []
  bestSBSs ← []
  $i \leftarrow 1$
  **while** $i \leq$ nIters **do**
    $S_{rand}$ ← knownSigs + $i$ randomly initiated sigs
    $A_{rand}$ ← randomly initiated activity matrix
    $A, S \leftarrow$ NMF($M, A_{rand}, S_{rand}$)
    banned ← banned + $s \in$ knownSigs that don't pass criteria
    knownSigs ← knownSigs - banned
    **if** $i == 1$ **then**
      novelSigs ← novelSigs + signatures that pass criteria and are not similar to any refSigs
    **end if**
    $\hat{S}$ ← signatures in $S$ that pass criteria and are not in knownSigs or banned
    bestSBSs ← bestSBSs + getBestSBSs(cosineSimilarity($\hat{S}$, refSigs))
    **if** at least one signature is in bestSBSs more than once **then**
      **for** $s \in$ bestSBSs more than once **do**
        knownSigs ← knownSigs + refSig with maximum cosine similarity to $s$
      **end for**
      $i \leftarrow 1$
    **else**
      $i \leftarrow i + 1$
    **end if**
  **end while**
  runSigs ← runSigs + knownSigs
**end for**
finalSigs ← signatures that appear in runSigs at least 0.8*nRuns times
$A_{rand}$ ← randomly initiated activity matrix
$A \leftarrow$ NMF($M, A_{rand}$,finalSigs)
novelClusterSigs ← median sig for each cluster(kmeans(novelSigs))
$A_1 \leftarrow$ NMF($M, A_{rand}$, (finalSigs + novelClusterSigs)
**Return:** finalSigs, $A$, novelClusterSigs, $A_1$

---

**Algorithm 2** getBestSBSs

**Input:** cosineSimilarity, $\hat{S}$, threshold, refSigs

similarSBSs ← []

**for** $s \in \hat{S}$ **do**

For each of the top 3 signatures in refSigs that have the highest cosineSimilarity to $s$, if their similarity ≥ threshold, append them to similarSBSs

**end for**

**Return:** similarSBSs

### 4 Hyperparameter Tuning

Since ReDeNovo allows users to adjust multiple parameters, we assessed how sensitive the results are to different parameter choices. We performed hyperparameter tuning to evaluate the sensitivity of the results to different parameter choices. Using synthetic data, we repeatedly ran the tool while systematically varying key hyperparameters (see Supplementary Table 6). Overall, the results indicate that ReDeNovo is robust to variations in hyperparameter settings.

While the tool is user-friendly and provides flexibility in parameter selection, we recommend adjusting parameters thoughtfully. For instance, we define a signature as valid if it appeared at least once, corresponding to the default value of the "–consno" parameter set to 1. Increasing this threshold up to 5 did not lead to any observable change in the results. While users may prefer slightly more conservative settings (e.g., values of 2 or 3), higher values are not recommended, as they may begin to degrade signal quality. As another example, thresholds for parameters controlling signature detection and exposure filtering must be chosen within a meaningful range. In our setup, increasing "thr1" beyond 0.10 and "exposure-thr-1" beyond 0.15 led to a drop in performance, since some exposures in our data occur only in subsets of the cohort. For such data, higher exposure thresholds should therefore be avoided. Likewise, if the dataset has low-magnitude exposures, "exposure-thr-2" should also remain relatively low to prevent loss of meaningful signal.

When we evaluated the impact of the number of iterations per run and the number of runs per experimental setup, no noticeable differences were observed; however, we recommend keeping both parameters sufficiently high (e.g., around 10) to achieve more consistent and stable results.

Below is a list of all parameters used by the tool and their definitions:

**User-defined parameters:**

- –consno: Minimum number of times a signature must be selected to be included in the inferred signature set. Default: 1
- -i or –numiters: Maximum number of iterations allowed while attempting to add new fixed signature (patience for novel signature). Default: 10
- -n or –numruns: Number of runs to repeat. Default: 10
- –thr1: Minimum fraction of patients with exposure >= thr1 required for a signature to be considered present. Default: 0.1
- –thr2: Minimum cosine similarity to match a signature with a known COSMIC signature and include in the inferred set. Default: 0.70
- –thr3: Minimum exposure weight for a signature to contribute to the final exposure profile. Default: 0.70 - present in 7 runs out of 10
- –thr4: Minimum cosine similarity to consider a signature as known and exclude it from novel candidate detection. Default: 0.80
- –thr5: Minimum fraction of the cohort with nonzero exposure required for a signature to be considered present. Default: 0.1
- –exposure-thr1: Minimum patient-wise normalized exposure required for a signature to be considered present. Default: 0.05
- –exposure-thr2: Minimum raw exposure required for a signature to be considered present. Default: 1
- -E or –exclude: List of SBS values to exclude from COSMIC, e.g. [”SBS1”, ”SBS5”]. Default: []

**Parameters about the database of catalogue signatures:**

- -g or –genome: Genome version, either 37 or 38. Default: 38.

- -w or –whole: Sequencing platform for the data and COSMIC catalogue. ”WGS” for Whole Genome Sequencing, ”WES” for Whole Exome Sequencing. Default: ”WGS”
- –cosmic-version: COSMIC version [”3.4”, ”3.3”, ”3.2”, ”3.1”, ”3”, ”2”, or ”1”]. Default: ”3.4”
- –manual-cosmic: Whether use manual COSMIC [user-provided COSMIC.txt in the input folder] (True or False). Default: False
- –manual-cosmic-file: Path to the file containing reference signatures. Used only if –manual-cosmic is set. Default: None

**Input file-related parameters:**

- -d or –delimiter: The delimiter used to separate the column in the input matrices. This delimiter will also be used for the output files. Default: '\t'
- –has-header-and-index: Whether the file has row names and column headers (True or False). Default: False
- –add-novel-signatures: Whether evaluate with novel signatures in the file provided (True or False). Used only if -N is set as 0. Default: False
- –check-novel: Whether check the novel signature or not (True or False). Default: False
- –novel-signatures-file: Path to the file containing novel signatures. Used only if –add-novel-signatures is set. Default: None

**Output-related parameters:**

- -O or –out: Path to output folder. Folder will be created if it does not not exist. If omitted, the current folder will be used as the output directory. Existing files will be overwritten.

[43] Aaron Chevalier, Tao Guo, Natasha Q Gurevich, Jingwen Xu, Masanao Yajima, and Joshua D Campbell. Characterization of highly active mutational signatures in tumors from a large chinese population. *medRxiv*, 2023.

[44] Hu Jin, Doga C Gulhan, Benedikt Geiger, Daniel Ben-Isvy, David Geng, Viktor Ljungström, and Peter J Park. Accurate and sensitive mutational signature analysis with musical. *Nature Genetics*, 56(3):541–552, 2024.

[45] Nicholas J Higham. Computing a nearest symmetric positive semidefinite matrix. *Linear algebra and its applications*, 103:103–118, 1988.

[46] Martín Abadi, Ashish Agarwal, Paul Barham, Eugene Brevdo, Zhifeng Chen, Craig Citro, Greg S Corrado, Andy Davis, Jeffrey Dean, Matthieu Devin, et al. Tensorflow: Large-scale machine learning on heterogeneous distributed systems. *arXiv preprint arXiv:1603.04467*, 2016.

[47] Diederik P Kingma. Adam: A method for stochastic optimization. *arXiv preprint arXiv:1412.6980*, 2014.
